# Ebselen protects XPC deficient cells through a potentially mitohormetic mechanism

**DOI:** 10.1101/2025.04.22.650012

**Authors:** Thiago S. Freire, Milena S. Martins, Neiliane Sima, Nadja C. de Souza-Pinto

## Abstract

Xeroderma pigmentosum group C fiborblasts (XP-C) are characterized by chronic redox imbalance and elevated H□O□levels, making them a good model for testing compounds with antioxidant potential for therapeutic purposes. Here, we investigated the effects of ebselen, a compound with glutathione peroxidase (GPx) mimetic activity, in the XP-C model. We found that ebselen behaves as a hormetic compound, protecting cells against H_2_O_2_-induced cytotoxicity at low doses but potentiating the cytotoxic effect at higher doses. Accordingly, when administered chronically, ebselen significantly reduces H□O□production and p53 levels. However, acute treatment with ebselen causes a reduction in O_2_ consumption (OCR) and extracellular acidification rate (ECAR), indicative of decreased mitochondrial function and metabolic activity. In addition, acute ebselen treatment causes a reduction in the GSH/GSSG ratio and an increase in NRF-2 expression, suggesting that ebselen induces redox stress that triggers an adaptive response, characterizing a possible mitohormetic effect. The reduction in the GSH/GSSG ratio appears to be the initial trigger after acute treatment with ebselen, since concomitant treatment with NAC prevents the reduction in OCR, ECAR and NRF-2 activation, in addition to protecting XP-C cells against lethal doses of ebselen.

**Highlights:**

- Chronic ebselen treatment reduces H_2_O_2_ levels, lowers p53 expression and protects XP-C cells against oxidative insults.
- Initially, ebselen causes mitochondrial stress, which is followed by cellular adaptation and protection against oxidative stress.
- Ebselen acts as a hormetic drug, likely through a mitohormetic mechanism.

## Introduction

Evolution favors the survival of organisms who better adapt to different stresses [1]. Thus, living beings have evolved stress response mechanisms that are engaged in a coordinated manner according to the survival needs, including DNA repair to ensure genomic stability [2]. Defects in these pathways can lead to human diseases [3], like defects in the nucleotide excision repair pathway (NER), which recognizes helix distorting damages. If not repaired, these lesions can lead to accumulation of mutations and cancer, as observed in xeroderma pigmentosum (XP) patients [4].

The XPC protein, mutated in XP complementation group C, participates in damage recognition in the NER pathway, which explains the high incidence of skin cancers in XP-C patients [5]. However, *in vitro* and *in vivo* models, and epidemiological data suggest that XPC deficiency is also associated with internal cancers, for which exposure to solar radiation is not relevant, implicating XPC in other biological roles than repair of UV-induced lesions via the NER pathway [6]. In fact, our group showed that XP-C cells display changes in expression and function of mitochondrial respiratory complexes and mitochondrial redox imbalance [7]. In addition, we showed that modulating H_2_O_2_ levels with N-acetylcysteine (NAC) reverses some of the phenotypes in the XPC-deficient cells, suggesting that the redox system plays an important role in this phenotype [8].

As H_2_O_2_ is likely the most relevant reactive oxygen species for signaling purposes, its levels are tightly controlled by systems that generate and degrade H_2_O_2_ according to signaling needs [9]. H_2_O_2_ acts mainly by oxidizing cysteine residues in target proteins, such that the reversible oxidation of thiols can act as redox switches, activating or deactivating enzymes. However, if H_2_O_2_ levels increase uncontrollably there may be additional oxidations that become irreversible, leading to damage to proteins, lipids and DNA [10]. Thus, the fine tuning of H_2_O_2_ generating and removing systems is extremely important. The mitochondrial electron transport chain plays a central role in generating H_2_O_2_, while the glutathione (GSH)/glutathione reductase (GR)/glutathione peroxidase (GPx) system constitutes one of the main H_2_O_2_ removing system [11]. In the XPC model, mitochondrial H_2_O_2_ generation is elevated, while GPx activity is decreased [7], thus favoring a dysfunction in the redox signaling pathways, which can explain the oxidative damage detected in these cells.

To test the hypothesis that preventing or reversing the redox imbalance seen in XP-C cells may prevent the dysfunctional phenotypes observed, we investigated the effect of ebselen, a selenium-containing small molecule developed as a GPx mimetic. Ebselen peroxidase activity has already been validated *in vitro* and *in vivo*, including in several clinical studies in humans, including for ischemic stroke [12]. Moreover, FDA has recently issued ebselen fast-track status for treating Meniere’s disease. Thus, ebselen has a clear potential as a pharmacological intervention to target redox dysfunction in disease models where redox imbalance may be relevant, such as XP-C.

## Materials and methods

Cell lines: The human SV40-transformed fibroblast cell lines XP4-PA (XP) carrying the dinucleotide deletion n.1747_1748delTG in exon 9 (formerly c.1643-1644delTG), and the corrected XP4-PA-CR cell line, complemented with the wild-type *XPC* gene [13], were kindly provided by Dr Carlos F. M. Menck, ICB, University of São Paulo. The cell lines were periodically checked for XPC expression by western blot (see Supplementary Figure S1). All cell lines were maintained in Dulbecco’s modified Eagle’s medium/high glucose (DMEM) supplemented with 10% fetal bovine serum (FBS), penicillin 100 IU/ml and streptomycin 100 μg/ml (pen-strep), at 37°C in a humidified atmosphere with 5% CO_2_. Cultures were routinely subcultured by trypsinization. For EB (Sigma– Aldrich) treatment, the cultures were maintained in DMEM supplemented with EB for two weeks for the chronic treatment, or for 24 hours for acute treatments. For the NAC (Sigma–Aldrich) treatment, the cultures were maintained in DMEM supplemented with 5mM of NAC for 24 hours before the experiments.

### Western blotting and in-cell western

whole cell extracts were prepared as described in [8] and quantified using the BCA assay (Pierce). Extracts were separated by SDS–PAGE in 12% denaturing gels and probed using antibodies against NRF-2, XPC, histone H3 (Abcam), p53 and β-actin (loading control; Sigma–Aldrich), using previously determined dilutions. For in-cell western, 4×10^4^ cells/well were seed in a 96-well plate and treated as described. After 24 hours cells were fixed with 50 µL/well 100% ethanol. Cells were blocked with BSA 1% for 1 hour at 37°C, followed by incubation with anti-p53 and anti-histone H3 for 24 hours at 4°C. Wells were washed 3x in PBS for 5 minutes each, added secondary antibody and incubated one hour at room temperature, washed three times in PBS for 5 minutes each and the plate left to dry. Fluorescence was measured using the Odyssey® XF Imaging System.

### Measurement of hydrogen peroxide levels

H_2_O_2_ formation was measured spectrophotometrically, in a SpectraMax 190 (Molecular Devices©), using Amplex® Red (Invitrogen) in the presence of 0.1 U/ml horseradish peroxidase (Sigma–Aldrich), as described in [7].

### Clonogenic assay

10^3^ cells/well were plated in 6-or 24-well plates and incubated for 24 h. Cells were washed 2x with phosphate-buffered saline (PBS) and incubated with H_2_O_2_ in DMEM without FBS for 30 min. After that, H_2_O_2_ was washed out with PBS and cells were incubated in complete medium for 6–7 days, until visible colonies were detected. Colonies were washed in ice-cold PBS, fixed with ethanol and stained with 1% (w/v) crystal violet solution. Survival rate was calculated as the ratio between the number of colonies in treated over non-treated conditions.

### Determination of O_2_ consumption

OCR and ECAR were measured using the XFe24 Extracellular Flux Analyzer (Agilent). Approximately 4□×□10^4^ cells/well were seeded in XFe24 V7 microplates and maintained at standard culture conditions. After 24□h, DMEM was replaced with 675 µL of DMEM without HCO_3_Na and cells were incubated in a humidified CO_2_-free incubator at 37□°C for 1□h. Respiratory parameters were analyzed following the manufacturer’s protocol for the Seahorse XF Cell Mito Stress Test Kit, with sequential additions of 1□*µ*M oligomycin, 200□nM FCCP and 1□*µ*M rotenone plus antimycin A, as described in [7].

### Measurement of GSH/GSSG ratio

GSH/GSSG ratio was determined using the GSH/GSSG-Glo™ Assay kit (Promega). Ten thousand cells/well were seed in a 96-well white plate. After 24 hours cells were treated with EB 5 µM for 4 hours; control wells received only DMEM. The medium was removed, and the samples were processed following the manufacturer’s protocol.

### Statistical analyses

results are shown as mean ± standard deviation (SD) of at least 3 independent experiments. ANOVA with Tukey regression or Sidak’s multiple comparisons test were used to compare differences of more than two groups, while Student’s t test was used to compare differences between two groups, considering significant p<0.05(*), p<0.01(**), p<0.005(***), p<0.001(****).

## Results

### Ebselen functions as a hormetic compound in XPC cells

We determined concentration range for XP-C cells incubated with ebselen (EB) for 24 h, using the XTT reduction as cell viability parameter (Figure 1A). Concentrations bellow 10 µM are well tolerated by XP-C cells, while concentrations above that induce dose-dependent, statistically significant cytotoxicity. Thus, further experiments were performed using EB concentrations bellow 10 µM, consistent with the concentration range used in the literature.

**Figure 1.**
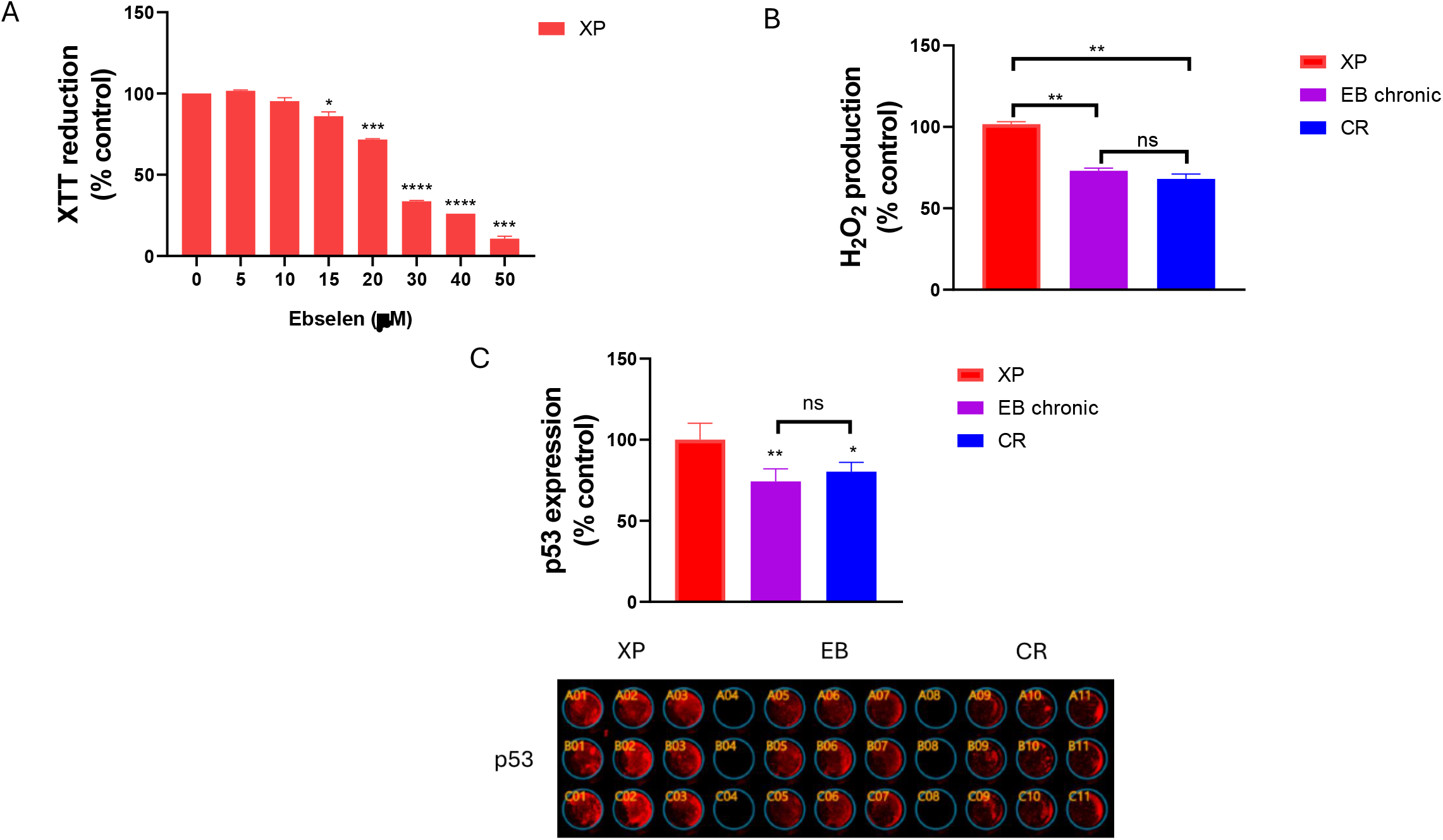
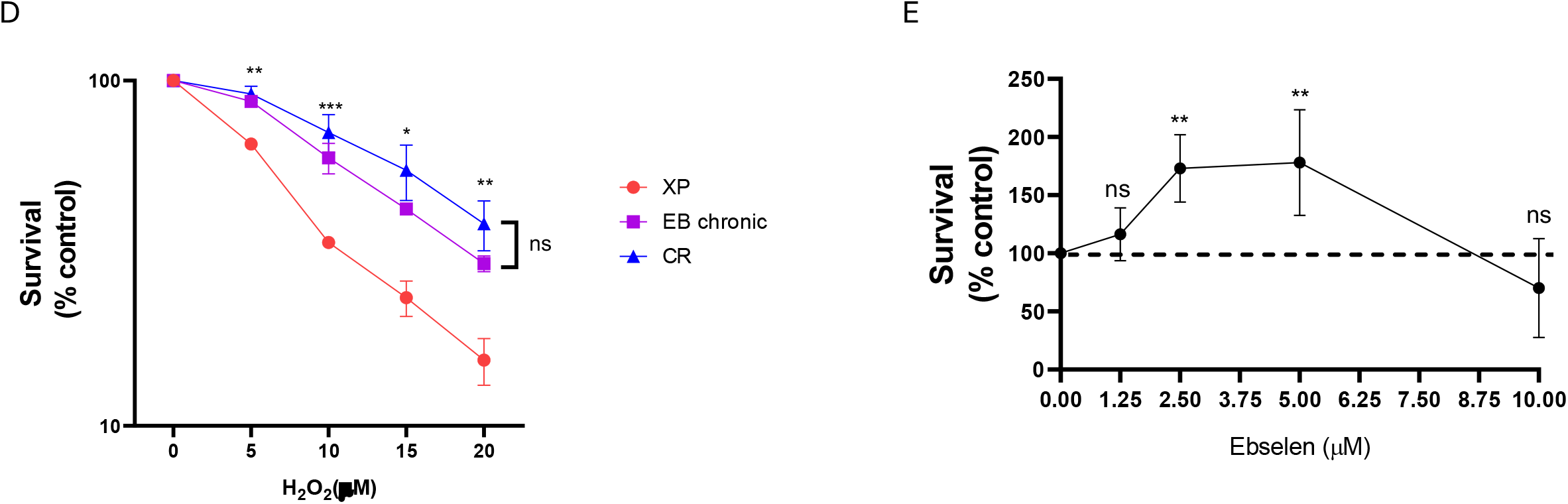
In all panels, **XP** refers to XP-C (XP4-PA) cells, **EB chronic** to XP-C treated with 5.0 µM ebselen every 48 hours for two weeks, and **CR** to XP-C cells corrected with the wild-type *Xpc* gene. (**A**) XP4-PA cells were treated with increasing concentrations of ebselen for 24 h, and cell density was determined by XTT assay. (**B**) H_2_O_2_ production was measured by Amplex Red assay. (**C**) Relative expression of p53 determined by In-Cell Western assay; the upper panel shows the quantification of 3 independent experiments and the bottom panel shows a representative image of the assay. (**D**) Cells were treated with increasing H_2_O_2_ concentrations and cell survival was determined by clonogenic assay. (**E**) Hormetic curve for increasing ebselen concentrations for XP-C cells treated with 10 µM H_2_O_2_, determined by clonogenic assay. In all panels, p<0.05(*), p<0.01(**), p<0.005(***), p<0.001(****).

Most EB biological effects are attributed to its GPx mimetic activity. Since H_2_O_2_ levels are elevated in XP-C cells [8], we investigate whether pre-treating XP-C cells with EB could modulate H_2_O_2_ levels. We used the same treatment protocol (two-week treatment, with fresh medium replaced every 48 h) used for N-acetyl cysteine (NAC), which showed effective in preventing H_2_O_2_ accumulation in the XP-C model [8]. Chronic treatment with 5 *μ*M EB significantly reduced H_2_O_2_ in XP-C cells, to levels equivalent to those detected in the wild-type corrected XP-C (CR) (Figure 1B), highlighting EB’s ability to modulate H_2_O_2_ in the XP-C cells.

We also showed that increased H_2_O_2_ levels correlated with p53 stabilization in XP-C cells [8]. Here, we found that EB treatment also reduces p53 levels to those found in CR cells (Figure 1C). In addition, chronic EB treatment restored resistance of XP-C to exogenously added H_2_O_2_ (Figure 1D), likely due to the reduction in endogenous H_2_O_2_ and p53 levels.

Since EB showed cytotoxicity at high acute concentrations and protection at low chronic concentrations, we speculated that EB acted in a hormetic fashion. To test that, we measured the effect of H_2_O_2_ on XP-C cells pre-treated with EB and found that pre-treating with 2.5-5.0 µM EB increased cell survival after H_2_O_2_, when compared with cells not treated with EB. Conversely, pre-incubation with 10 µM EB renders cells more sensitive to H_2_O_2_, displaying a typical hormetic profile (Figure 1E).

### Mitochondria are an early EB cellular target

Some hormetic compounds, like 2-deoxy-D-glucose and metformin, exert their effects mostly or partially through mitochondria [14]. We tested whether EB had direct effects on mitochondrial respiration, measuring O_2_ consumption rate (OCR) and extracellular acidification rate (ECAR) in cells treated with EB either for 24 h (acute) or 2 weeks (chronic). Acute EB treatment significantly reduced OCR in XP-C cells, particularly basal and maximum respiration (Fig 2A, lower panel). Lower mitochondrial respiration was not compensated by increased glycolytic activity, as indicated by significantly lower ECAR in the same conditions (Fig. 2B). On the other hand, cells subjected to the chronic protocol presented OCR and ECAR like the untreated control. These results suggest that mitochondrial dysfunction and metabolic stress are early events in the EB hormetic mechanism. Given that EB chronically treated cells are more resistant to H_2_O_2_-induced oxidative stress, this result suggests that EB could act as a mitohormetic compound.

**Figure 2.**
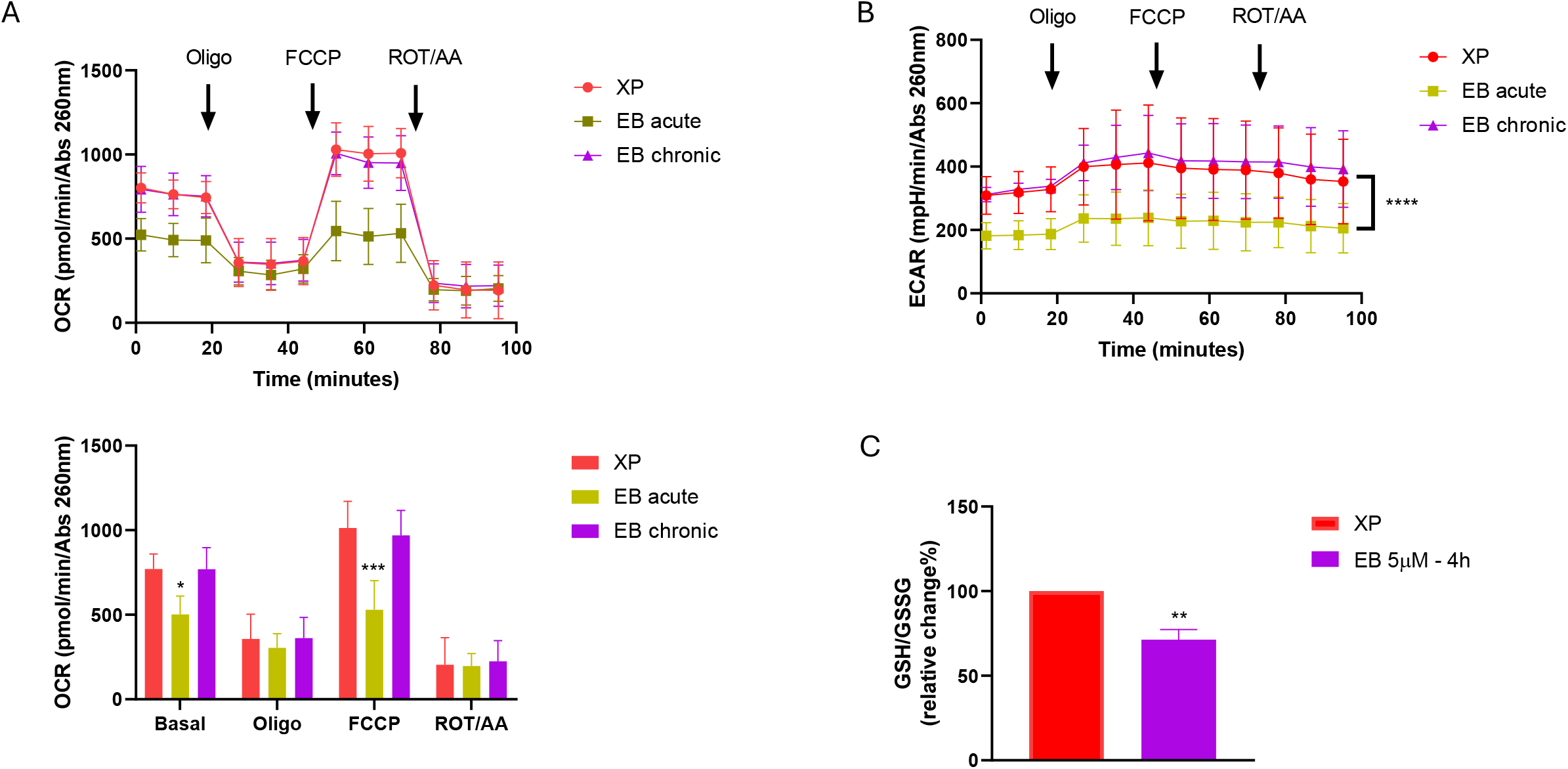
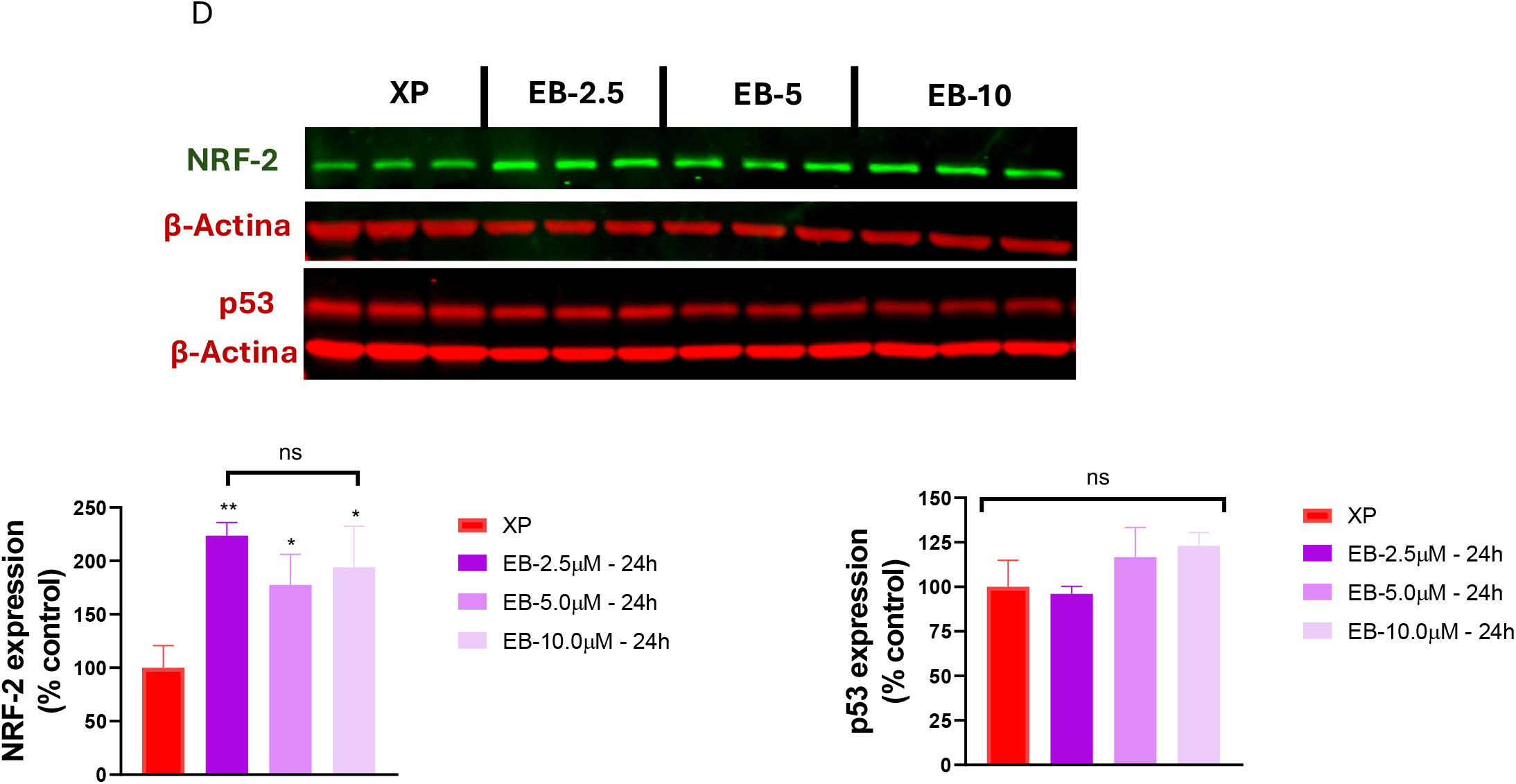
In all panels, **XP** refers to XP-C (XP4-PA) cells, **EB acute** refers to XP-C treated with 5.0 µM ebselen 24 hours before the experiment, and **EB chronic** to XP-C treated with 5.0 µM ebselen every 48 hours for two weeks. (**A**) O_2_ consumption rate (OCR) determined by Seahorse platform, after additions of oligomycin (1□*µ*M), FCCP (200□nM) and rotenone/antimycin A (1□*µ*M each). The top panel shows the data in graph format while the bottom panel shows quantification of the individual parameters. (**B**) Extracellular acidification rate (ECAR) in the same conditions as above. (**C**) GSH/GSSG ratio was measured in control XP-C cells (XP) or treated with 5.0 µM ebselen for 4 h before the experiment (EB 5μM -4h). NRF2 and p53 levels were measured, by western bloting, in control XP-C cells or treated with increasing concentrations (2.5-10 µM) ebselen for 24 h (**D**) or 4 days (**E**). In panels D & E, the top figure shows a representative image of the western blot and the bottom graphs show the quantification normalized to β-actin levels. Results shown are average ± SD of 3 independent experiments, performed in quadruplicate, p<0.05(*), p<0.01(**), p<0.005(***), p<0.001(****).

### EB exerts its mitohormetic function though redox modulation

Since EB is a GPx mimetic, we hypothesized that EB accelerated GSH oxidation reducing the GSH/GSSG ratio, and that the stress resistance seen after chronic EB treatment was a consequence of these initial effects. We tested this hypothesis and found that 4 h treatment with 5 µM EB significantly reduced GSH/GSSG ratio in XP-C cells (Figure 2C).

Consistent with an increased redox imbalanced induced by EB in the initial times, NRF2, an important transcription factors for redox-induced adaptive gene expression, levels were significantly upregulated after 24 h EB treatment (Figure 2D, left panel). Interestingly, there was no significant change in p53 levels (Figure 2D, right panel).

The hormetic hypothesis presupposes that the initial stress caused by EB leads to an adaptive response that results in better antioxidant defense. After 4 days treatment with EB both NRF-2 and p53 levels reduced significantly, indicative of a reorganization of the redox homeostasis (Figure 2E). We then tested the effects of N-acetyl cysteine (NAC), which we previously shown to upregulate GSH levels in XP-C cells [8]. NAC pretreatment protected XP-C cells against dose-dependent EB acute toxicity, even at 20 µM EB, where there is practically no cell survival in absence of NAC (Figure 3A), indicating that EB’s initial cytotoxicity is directly linked to GSH depletion.

**Figure 3.**
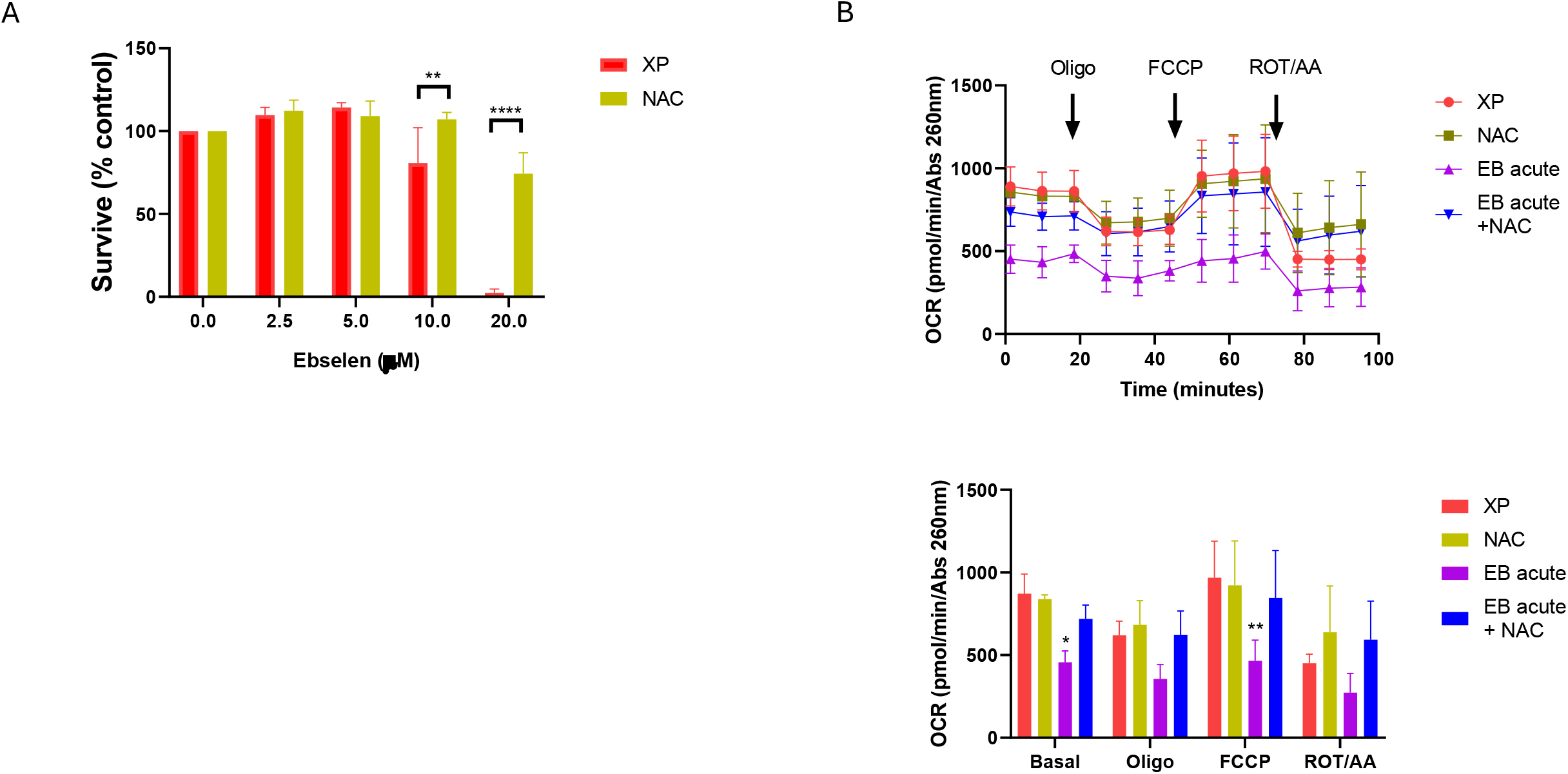
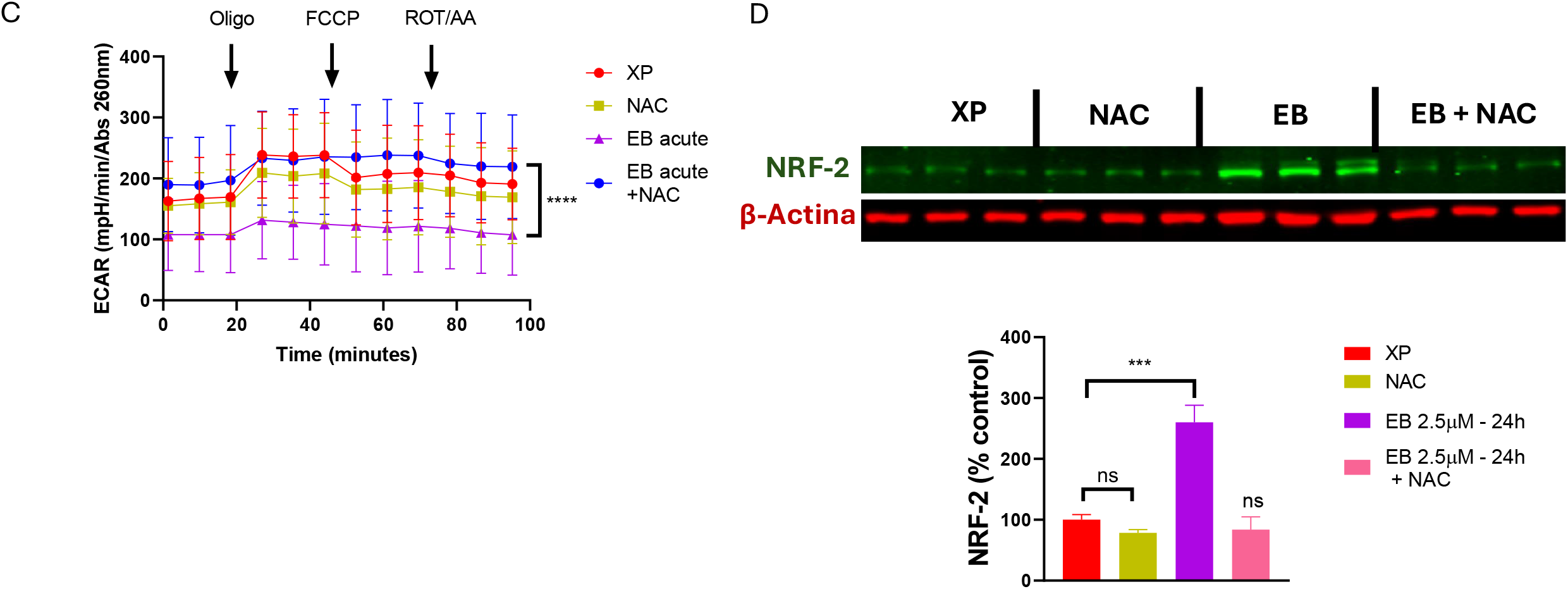
(**A**) XP-C cells pre-treated with DMEM alone (XP) or DMEM containing 5 mM NAC were treated with increasing concentrations of ebselen form 24 h, and the relative survival was determined by clonogenic assay. XPC-cells pre-incubated with DMEM alone (XP) or 5 mM NAC (NAC) were treated with 5.0 µM ebselen 24 hours before the experiment (EB acute and EB acute + NAC, respectively). After the treatment, OCR (**B**) and ECAR (**C**) were determined, as previously. In (**D**) XPC-cells pre-incubated with DMEM alone (XP) or 5 mM NAC (NAC) were treated with 2.5 µM ebselen for 24 hours and the relative expression of NRF-2 was determined by western blotting. Results are average ± SD of 3 independent experiments. In all panels, p<0.05(*), p<0.01(**), p<0.005(***), p<0.001(****).

Interestingly, EB-induced decreases in basal and maximum OCR (Figure 3B), and in ECAR (Figure 3C) were prevented by pre-incubating with NAC. And in agreement with the hypothesis that the reduction in the GSH/GSSG ratio and, possibly, mitochondrial and metabolic stress are the triggers for activating NRF-2, pretreatment with NAC completely blocked acute EB-induced NRF-2 expression upregulation (Figure 3D).

## Discussion

EB is a small selenium-containing molecule with GPx-mimetic catalytic activity, i.e., it oxidizes GSH to GSSG and reduces H_2_O_2_ to H_2_O. There is sufficient literature data and human clinical trials demonstrating its effectiveness as an antioxidant, anti-inflammatory and neuroprotective agent [15]. Here, we find that EB concentrations of up to 10 µM are not cytotoxic to XP-C cells (Figure 1A), with the assays used here. Thus, we chose to use values within this range (0-10 µM), which are well within the range used in the literature.

Our results support the hypothesis that EB acts as an antioxidant. Chronic treatment with 5.0 µM EB for two weeks, with fresh doses at 48-hour intervals, reduced H_2_O_2_ (Figure 1B) and p53 levels (Figure 1C). These results agree with our previous results that H_2_O_2_ modulates p53 levels in XP-C cell, such that treatments that lower the former should, in turn, reduce the latter, as we see here [8].

Several studies have shown that XPC-deficient models are more sensitive to conditions that cause oxidative stress [7]. This is mainly because XP-C cells appear to produce more oxidants while being deficient in some antioxidant enzymes [7]. Interestingly, chronic EB lowers H_2_O_2_ levels and protects XP-C cells against exogenously added H_2_O_2_ (Figure 1D). The relevance of these results lies in the fact that a decrease in oxidant production and p53 levels, and an increase in antioxidant defenses are conditions known to protect cells from the carcinogenic process, as demonstrated by results showing that reducing redox imbalance in XP-C can reduce lung tumors [16].

Literature data shows that high doses of EB disrupt mitochondrial function [17]. We hypothesized that the chronic treatment allowed enough time for transcriptional rearrangements that lead to adaptation. Thus, to understand the initial effects induced by EB, we investigated the acute effects in XP-C cells.

Treatment with 5 µM EB for 24 hours significantly reduced OCR (Figure 2A) and ECAR (Figure 2B) in XP-C cells, suggesting that initially EB causes a metabolic stress. On the other hand, OCR and ECAR of cells treated with EB chronically are not significantly different from untreated control (Figures 2A/2B, respectively). Thus, we hypothesize that EB causes an initial mitochondrial and metabolic stress that activates adaptive responses that result in a more robust antioxidant profile. The phenomenon in which an agent induces an initial stress capable of activating cellular adaptation systems is called hormesis [18], or mitohormesis when the first target are mitochondria [14]. Our results suggest that EB potentially exerts some of its effects via mitohormesis. Other molecules are proposed to act though mitohormesis, like metformin, a drug widely used to treat type 2 diabetes, and known to protect against cardiovascular diseases, cancers and dementia via a mechanism seemly unrelated to its role in regulating glycemic levels [19].

Our results support the hypothesis that EB acts as a hormetic compound (Figure 1E). Since EB is a GPx mimetic, we hypothesized that the initial trigger would be a reduction in the GSH/GSSG ratio due to increased GSH oxidation. In fact, acute EB treatment reduces GSH/GSSG ratio (Figure 2C), which was prevented by NAC, which protects against lethal dose of EB (Figure 3A). Furthermore, acute EB treatment upregulated NRF-2 expression, a pivotal factor in the transcriptional response against oxidative insults, with no apparent effect on p53 levels (Figure 2D). On the other hand, after 4 days, both NRF-2 and p53 levels were markedly reduced, suggesting that the initial stress was resolved (Figure 2E).

The GSH/GPx system that oxidizes GSH to GSSG is in balance with the NADPH/GR system that reduces GSSG to GSH, ensuring that the GSH/GSSG ratio remains at levels compatible with the cell’s redox homeostasis [20]. EB increases GSH consumption, however with no immediate effect on the activity of the NADPH/GR system. As a result, at the earlier times in presence of EB there is a reduction in the GSH/GSSG ratio, disrupting redox balance (Figure 2C). This occurs because the GSH/GSSG ratio affects, directly and indirectly, the GRx (glutaredoxin) and TRx (thioredoxin) systems, which ensure the redox homeostasis of cysteines in proteins, consequently controlling and regulating their activity and function [21]. We propose that the initial reduction in the GSH/GSSG ratio would be the trigger that led to reduction in mitochondrial respiration. In agreement with this, studies using selenium-(including EB) and tellurium-containing organic compounds in isolated mitochondria, showed that both reduced respiratory complexes I and II activities, and this was probably caused by cysteines oxidation, given that concomitant treatment with GSH prevented this effect, but treatment with superoxide dismutase or catalase had no effect. These results suggested that the effect is caused by GSH depletion itself and not by an increase in H_2_O_2_ or O_2_ ^•-^ levels. [22]. We showed here that EB treatment reduced mitochondrial respiration in intact cells, and that co-treatment of EB with NAC, which maintains intracellular GSH concentration, prevented cytotoxicity at high EB concentrations (Figure 3A), decrease in OCR (Figure 3B) and in ECAR (Figure 3C). Furthermore, EB+NAC completely blocked NRF-2 upregulation (Figure 3D). Together, these results suggest that the reduction in GSH levels and mitochondrial and metabolic stress are the triggers for the activation of the adaptive response that begins after acute EB treatment.

NRF-2 is an important transcription factor that responds to oxidative and electrophilic insults. Under stress conditions, NRF-2 is stabilized, migrates to the nucleus and activates a set of genes responsible for maintaining cellular redox homeostasis. Among the responses mediated by NRF-2 are the transcription of genes of the GSH/GSSG system, such as GR enzyme and genes related to NADPH production. Thus, NRF-2 activation can restore homeostasis of the GSH/GSSG system [23]. Here we show that acute treatment with EB increases NRF-2 levels (Figure 2D), which would, in turn, induce expression of adaptive genes and increase antioxidant capacity. This response would decrease chronic redox stress and, subsequently, reduced expression of oxidative stress-responsive factors, p53 and NRF-2 (Figure 2E).

In conclusion, our mechanistic proposal is that EB treatment causes an initial GSH depletion that leads to cysteine oxidation in subunits of the electron transport chain complexes, causing reduced bioenergetic activity and metabolic stress. This initial stress is the event that activates NRF-2, which orchestrates the expression of antioxidant response genes, including GR, therefore, reestablishing the redox balance. In this scenario, GR and GPx activities would return to equilibrium, but with higher activities than those under basal conditions, resulting in decreased H_2_O_2_ levels and a more robust antioxidant system, as schematized in Figure 4.

**Figure 4.**
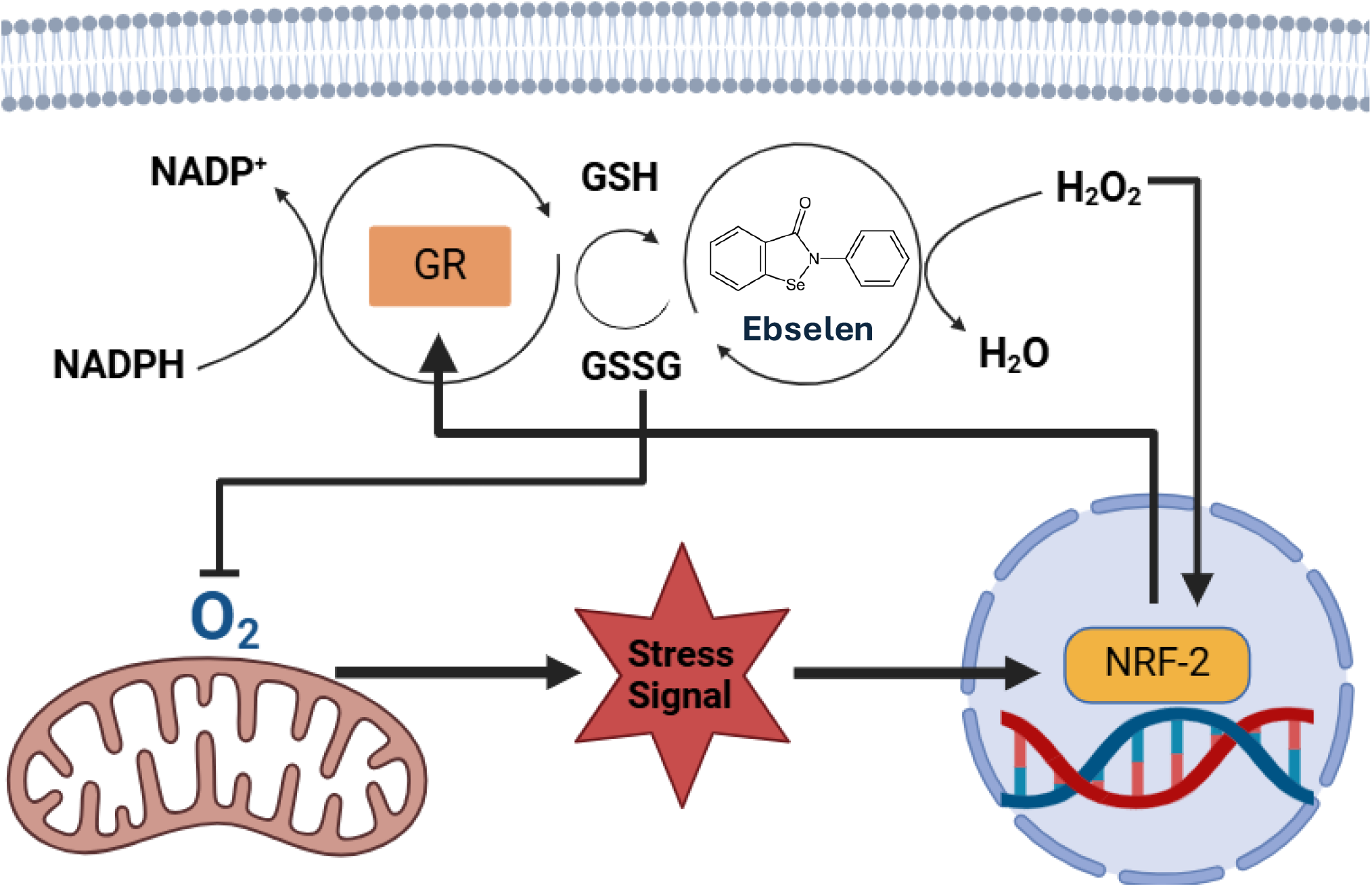
Mitohormetic mechanistic hypothesis. Initially, ebselen increases GSH oxidation, imposing on mitochondrial function to maintain the redox homeostasis. This creates a stress signal capable of inducing NRF-2expression, which in turn can activate the expression of genes that will reestablish the balance of the GSH/GSSG system, consequently increasing the efficiency of the antioxidant system against oxidative stress induced by H_2_O_2_.

## Abbreviations

EB: ebselen
XP-C: xeroderma pigmentosum complementation group C
OCR: oxygen consumption rate
ECAR: extracellular acidification rate
NRF-2: nuclear factor erythroid-2-related factor 2
GSH: reduced glutathione
GSSG: oxidized glutathione
NAC: N-acetylcysteine
PBS: phosphate-buffered saline.

## Acknowledgments

This work was supported by Fundação de Amparo à Pesquisa do Estado de São Paulo (FAPESP), grants # 2017/04372-0 and 2022/01155-7, and Programa Unificado de Bolsas da Universidade de São Paulo, project # 2023-5291. The authors thank Adriana M. P. Wendel for technical assistance.

