## Supplemental Figure 1 for "Ebselen protects XPC deficient cells through a potentially mitohormetic mechanism"

### Slide 1
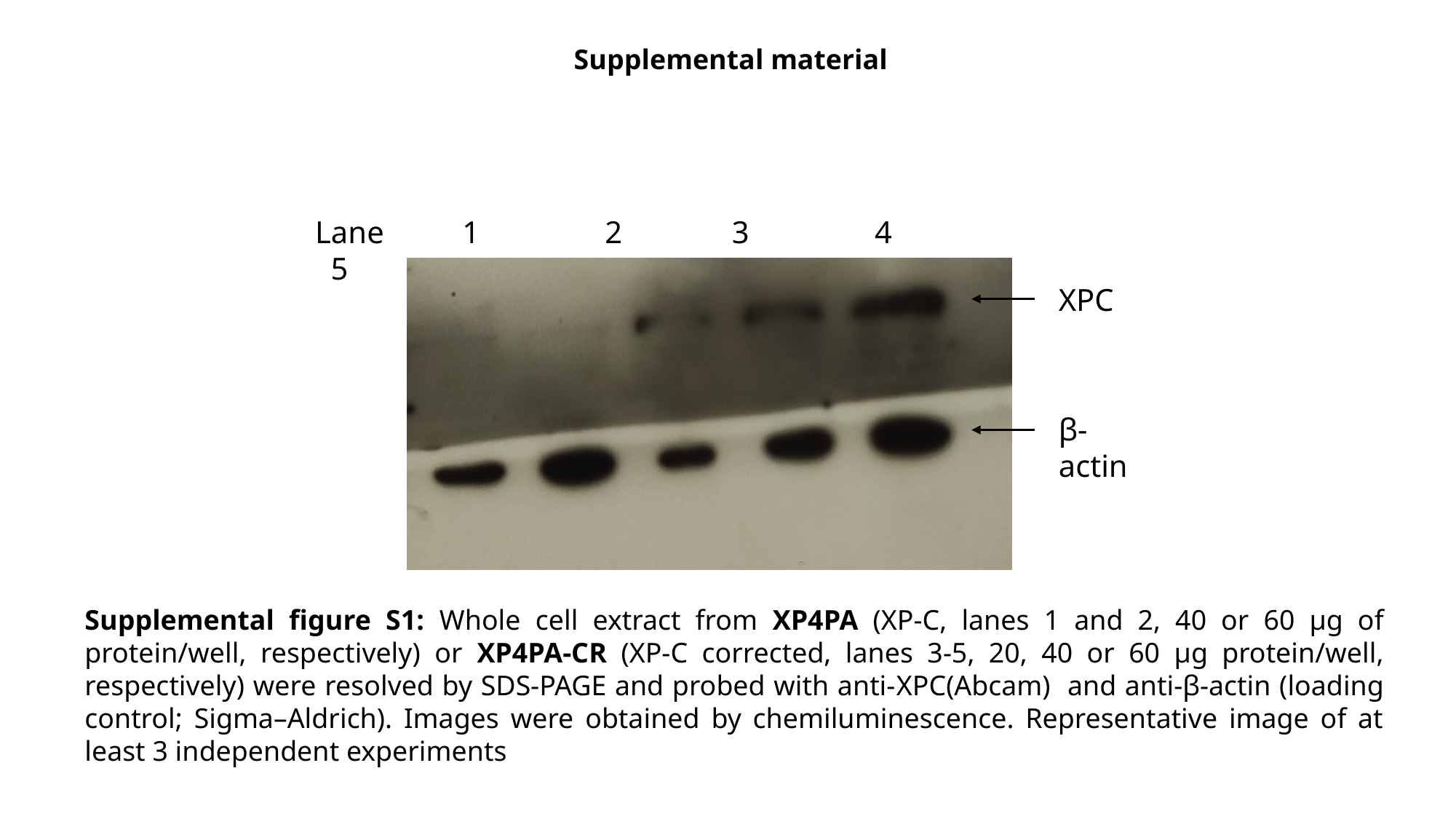

Supplemental material
Lane 1 2 3 4 5
XPC
β-actin
Supplemental figure S1: Whole cell extract from XP4PA (XP-C, lanes 1 and 2, 40 or 60 µg of protein/well, respectively) or XP4PA-CR (XP-C corrected, lanes 3-5, 20, 40 or 60 µg protein/well, respectively) were resolved by SDS-PAGE and probed with anti-XPC(Abcam) and anti-β-actin (loading control; Sigma–Aldrich). Images were obtained by chemiluminescence. Representative image of at least 3 independent experiments
